# *TML1* AND *TML2* SYNERGISTICALLY REGULATE NODULATION AND AFFECT ARBUSCULAR MYCORRHIZA IN *MEDICAGO TRUNCATULA*

**DOI:** 10.1101/2023.12.07.570674

**Authors:** Diptee Chaulagain, Elise Schnabel, Mikayla Kappes, Erica Xinlei Lin, Lena Maria Müller, Julia A. Frugoli

## Abstract

Two symbiotic processes, nodulation and arbuscular mycorrhiza, are primarily controlled by the plant’s need for nitrogen (N) and phosphorus (P), respectively. Autoregulation of Nodulation (AON) and Autoregulation of Mycorrhization (AOM) both negatively regulate their respective processes and share multiple components - plants that make too many nodules usually have higher AM fungal root colonization. The protein TML (TOO MUCH LOVE) was shown to function in roots to maintain susceptibly to rhizobial infection under low N conditions and control nodule number through AON in *Lotus japonicus*. *M. truncatula* has two sequence homologs: *Mt*TML1 and *Mt*TML2. We report the generation of stable single and double mutants harboring multiple allelic variations in *MtTML1* and *MtTML2* using CRISPR-Cas9 targeted mutagenesis and screening of a transposon mutagenesis library. Plants containing single mutations in *Mt*TML1 or *Mt*TML2 produced 2-3 times the nodules of wild-type plants whereas plants containing mutations in both genes displayed a synergistic effect, forming 20x more nodules compared to wild type plants. Examination of expression and heterozygote effects suggest genetic compensation may play a role in the observed synergy. Plants with mutations in both *TMLs* only showed mild increases in AM fungal root colonization at later timepoints in our experiments, suggesting these genes may also play a minor role in AM symbiosis regulation. The mutants created will be useful tools to dissect the mechanism of synergistic action of *Mt*TML1 and *Mt*TML2 in *M. truncatula* symbiosis with beneficial microbes.

## Introduction

Since plants are sessile and require nutrients from the soil, they have developed strategies to maximize survival when the supply of nutrients is variable (Oldroyd and Leyser, 2020). Legume plants can grow in nitrogen (N) poor soil using an additional strategy beyond those of most dicots: legumes can establish an endosymbiosis with Rhizobia bacteria, which allows the N fixed from the atmosphere by the bacteria to be shared with the plant. Rhizobia are harbored in specialized root organs called nodules. The establishment of the legume root nodule symbiosis is tightly regulated, including by species-specific signals from the rhizobia (lipo-chitooligosaccharides) and internal plant signaling related to the carbon and N status of the plant. Obtaining N through symbiosis is energetically costly to the plant, resulting in regulation based on both available soil N, which appears to use common signals among most plants (Oldroyd and Leyser, 2020), and on already established nodulation, which uses a pathway called Autoregulation of Nodulation (AON) to control the number of nodules that form on a legume (reviewed in (Ferguson et al., 2019; Roy et al., 2020; Chaulagain and Frugoli, 2021)).

The AON pathway negatively controls nodule number through systemic signal transduction and has been studied in many species; here we use *Medicago truncatula* nomenclature. Upon inoculation of *M. truncatula* with rhizobia, genes encoding the signaling peptides MtCLE12 and MtCLE13 are induced in the root (Mortier et al., 2010) through activation by NIN (Laffont et al., 2020). All CLE peptides undergo proteolytic processing from the original gene transcripts (see (Gao and Guo, 2012; Roy and Müller, 2022) for review) and MtCLE12 is additionally post translationally glycosylated by the MtRDN1 enzyme before being translocated to the shoot (Kassaw et al., 2017). In the shoot, these CLE peptides are ligands for the leucine-rich receptor-like kinase (LRR-RLK) *M. truncatula* SUNN (SUPERNUMERARY NODULES) (Mortier et al., 2012). SUNN, which is the *M. truncatula* ortholog of Arabidopsis CLAVATA1, forms a putative complex with the pseudo kinase CORYNE (CRN) and the receptor protein CLAVATA2 (CLV2) (Crook et al., 2016). MtCLE12,13/SUNN signaling results in increased transport of cytokinin and decreased expression the of shoot-to-root signal miR2111 (Tsikou et al., 2018; Okuma et al., 2020; Okuma and Kawaguchi, 2021).

Based on mutational analysis in *L. japonicus* which has only one *TML (TOO MUCH LOVE*) gene and RNAi in *M. truncatula* which has two, reduced levels of miR2111 in roots correspond with increased expression of the *TML* genes, which encode F-box proteins that negatively regulate nodule number (Magori et al., 2009; Tsikou et al., 2018; Gautrat et al., 2019). The downregulating effect of miR2111 on *TML* transcripts is postulated based on several lines of evidence: (1) the presence of predicted binding sites for miR2111 in the 5’ end of the *TML* gene (Gautrat et al., 2020), (2) the observation that a miR2111 target mimic increases the *TML* transcript levels in a *sunn* mutant (Gautrat et al., 2020), and (3) increasing miR2111 levels through overexpression in roots increases nodule number and decreases the transcript levels of *TML* at five days post inoculation (dpi) (Gautrat et al., 2021).

Like nodulation, plant interactions with arbuscular mycorrhiza (AM) fungi are also controlled by autoregulation and plant nutrient status. AM symbiosis occurs in most land plants, including 85-90% of angiosperms; unlike nodulation AM symbiosis is not restricted to legumes (Genre et al., 2020). The autoregulation and nutrient-regulated signaling pathways fine-tuning AM symbiosis share several, but not all, components and mechanisms with AON and N regulation of nodulation (Bashyal et al., 2023). In *M. truncatula*, autoregulation of mycorrhizal symbiosis (AOM) is mediated by the AM-induced peptide MtCLE53, which elicits a negative feedback loop that represses AM symbiosis in concert with the LRR-RLK SUNN (Müller et al., 2019; Karlo et al., 2020). Components of AOM are conserved beyond the legume clade: hyper-mycorrhizal mutants – characterized by elevated overall root length colonization and arbuscule numbers – in CLE peptides and orthologs of SUNN and CLV2 have been described not only in the legumes *M. truncatula*, soybean, pea, and *Lotus japonicus*, but also in the non-legumes *Brachypodium distachyon* and tomato (Morandi et al., 2000; Zakaria Solaiman et al., 2000; Meixner et al., 2005; Wang et al., 2018; Müller et al., 2019; Karlo et al., 2020; Wang et al., 2021). Notably, recent research revealed the existence of multiple parallel CLAVATA signaling pathways modulating AM symbiosis (Orosz et al., 2024; Wulf et al., 2024).

AM symbiosis is also regulated by plant P and N status. Accumulating evidence suggests that P- and N-induced CLE peptides contribute to nutrient regulation of AM symbiosis; although our knowledge is still fragmented, at least some of the nutrient-regulated CLE peptides require the same LRR-RLKs or other regulators of AOM and AON (Müller et al., 2019; Wang et al., 2021).

Thus far*, TML* is the only gene with a loss of function hypernodulation mutant phenotype reported to function downstream of the SUNN LRR-RLK in the AON pathway and was first described in *L. japonicus* (Magori et al., 2009). No mutation of a gene downstream of SUNN resulting in a hyper-mycorrhizal phenotype has been reported in AOM. The *L. japonicus tml* mutant plants form ∼8x more nodules spreading across larger area of root than wild type, a phenotype governed by root genotype (Magori et al., 2009). The gene encodes a protein containing a Nuclear Localization Signal (NLS), F-box domain, and kelch repeats (Takahara et al., 2013). Previous work demonstrated overexpression of *MtCLE12* and *MtCLE13* results in induction of *MtTML1* and *MtTML2* expression even in absence of rhizobia (Gautrat et al., 2019). In addition, MtCEP1 (*C-TERMINALLY ENCODED PEPTIDE1*) generated in response to low N, requires the MtCRA2 (*COMPACT ROOT ARCHITECTURE2*) receptor to upregulate miR2111 expression in the shoots and hence lower the transcript level of both *MtTML*s to maintain root competence to nodulation (Gautrat et al., 2020).

The presence of kelch repeats and F-box domains in TML proteins suggest TMLs could bind to each other, as well as target other proteins for proteasomal degradation, however biochemical studies to identify such targets have not occurred yet. Genetic studies in *M. truncatula* have been hindered by the lack of stable mutants and have been limited to RNA interference (RNAi) experiments and measuring transcript levels (Gautrat et al., 2019; Gautrat et al., 2020). To understand the role of two TML proteins in nodulation, mutant analysis is critical. We obtained a mutant in *MtTML2*, *tml2-1*, from a *Tnt1* transposon insertion mutant library (Tadege et al., 2008) and generated stable single and double mutants harboring multiple allelic variations in *MtTML1* and *MtTML2* using CRISPR-Cas9 targeted mutagenesis. Plants containing mutations in a single *MtTML* gene displayed two to three times as many nodules as wild type plants, however plants containing mutations in both genes displayed a synergistic effect, with up to twentyfold more nodules than the wild type plants. By contrast, AM fungal root colonization was only mildly enhanced in a *tml* double mutant; furthermore, this effect was dependent on the timepoint measured and the mutant allele. Taken together with observations on gene expression and other phenotypes reported below, our findings demonstrate the requirement of both *MtTML1* and *MtTML2* in controlling nodule number through synergistic signaling in nodulation with an additional minor role in AM symbiosis regulation under the experimental conditions tested.

## Materials and methods

### Plant Growth Conditions

Seeds were germinated as described previously (Schnabel et al., 2010). Briefly, seeds were scarified using concentrated sulfuric acid, imbibed in water, vernalized at 4°C for 2 days and allowed to germinate overnight at room temperature in dark. One day old seedlings were placed in an aeroponic chamber and grown at 21°C-25°C; 14h/10h light/dark cycle and inoculated as described in (Cai et al., 2023). To determine nodulation phenotypes no N was included in the media and plants were inoculated *Sinorhizobium meliloti* RM41 (Putnoky et al., 1990) four days post germination as indicated. Nodule count and root length measurements were performed 10 days post inoculation (dpi). For seed collection and genetic crosses, plants were grown in a greenhouse with supplemental light on a 14h/10h light/dark cycle at 21°C-25°C. For mycorrhizal assays, seed germination and inoculation with *Rhizophagus irregularis* was performed as previously described (Chaulagain et al., 2023). In brief, 3-day old seedlings were planted in Cone-tainers SC10R (Steuwe and Sons) and placed in a growth chamber with 16h/8h light/dark cycle at 22°C-24°C and 40% humidity. The growth mixture was 50% play sand (Quickrete) and 50% fine vermiculite (Ferry-Morse). All substrate components were washed in distilled water and autoclaved for 55 minutes before use. For colonization with *R. irregularis*, 250 spores (Premiertech, Canada) were placed 5 cm below the surface in a layer of fine sand. All plants were treated twice a week with 15ml of half-strength, low Pi Hoagland’s fertilizer (20µM phosphate) to aid in symbiosis initiation. Plants were harvested 4.5 or 6 weeks post planting.

### Tnt1 mutant screening and verification of insert

*Tnt1* insertion lines created by *Tnt1* mutagenesis of the R108 ecotype (Tadege et al., 2008) were screened electronically for insertions in *MtTML* genes. While no *MtTML1* insertion lines were identified, a pool of plants containing a mutation in the *MtTML2* gene in *M. truncatula* (NF0679) Noble Foundation Medicago Mutant Database now located at https://medicagomutant.dasnr.okstate.edu/mutant/index.php. Seeds from the NF0679 pool were grown in the greenhouse and a line containing the insertion in the second exon was identified by PCR using primers from Supplemental Table 2. The line was selfed to yield homozygotes, which were then backcrossed to the R108 wild type and re-isolated. The amplicons generated by primers 2189/1925 and 2690/3227 were sequenced to determine the exact position of the *Tnt1* insertion in relation to the gene. This line is called *tml2-1*.

### Constructs for CRISPR-Cas9 mutagenesis

Multi target constructs were designed for *MtTML1* alone and to target both *MtTML* genes in the same plant (Supplemental Figure 1). Two target sites per gene were chosen following the target selection criteria (Ma and Liu, 2016) Sites unique to the *MtTML* genes were selected to avoid off-target mutagenesis by comparison to the *M. truncatula* genome MtV4.0 (Tang et al., 2014). The binary vector pDIRECT_23C (Čermák et al., 2017) (Addgene) was used as Cas9 containing vector. The target and gRNA were cloned into the pDIRECT_23C vector and the finished construct created using Golden Gate assembly (Engler et al., 2008; Engler et al., 2009) following the protocol in (Čermák et al., 2017). The resulting vectors pDIRECT_23C+MtTML1 (containing two targets for *MtTML1*) and pDIRECT_23C+MtTML1/2 (containing two targets each for *MtTML1* and *MtTML2)* were then verified by sequencing. A verified clone in *E. coli* (Zymo 10B, Zymo Research) was used as plasmid source to move each construct into *Agrobacterium tumefaciens* EHA105 (Hood et al., 1993) for use in plant transformation.

### Generation of transgenic plants using CRISPR-Cas9 mutagenesis

To obtain alleles of *MtTML1*, additional alleles of *MtTML2* and plants containing mutations in both genes, CRISPR-Cas9 mutagenesis of embryonic callus tissue followed by regeneration of whole plants was performed in the R108 ecotype. Each construct was introduced with *Agrobacterium-*mediated transformation into root segments of 4-7 day old R108 seedlings and taken through callus and tissue culture following the protocol 2A of (Wen et al., 2019) with the following modifications for selection and rooting: the selection of phosphinothricin (ppt) resistant callus used 2mg/L ppt in the media and 2mg/L NAA was used in RCTM6 medium described in (Wen et al., 2019) for rooting. All plants originating from a callus were considered one independent line. The three most developed plants at the rooting medium stage for each line per callus were transferred into soil for acclimatization and grown to collect seeds.

### Transgenic verification, genotyping, and segregation of targeted mutagenesis in T1 generation

All T0 lines generated from tissue culture were screened by PCR for the presence of the transgene using Cas9 specific primers (primers 3081/3082; Supplemental Table 2), and lines testing positive were then screened by PCR for deletions using gene specific primers (Supplemental Table 2). Deletion was confirmed by size difference of an amplicon after gel electrophoresis. All the amplicons were sequenced using the same primers as for the PCR to determine the exact mutations.

All T0 lines containing mutations were allowed to set seeds. The T1 generation was screened for plants homozygous for a single allele by PCR. All amplicons were sequenced to confirm homozygosity and sequence. To generate the *tml1 tml2* double mutant transgenic lines, a T0 line carrying two mutant alleles for each gene was used. All alleles except *tml1-4* were backcrossed once to R018 and all alleles including the homozygous double mutants do not have detectable Cas9 when tested by PCR amplification.

### Real Time PCR Analysis of MtTML in mutant plants

RNA was extracted from 10 roots each of mutants and R108 (wild type -WT) harvested at 10 days post inoculation with RM41. Root tissue was frozen at −80°C until RNA extraction. Root tissue from six plant pooled together for each genotype was ground in liquid nitrogen and 100mg of tissue was used for RNA extraction. RNA was extracted using TRIzol® Reagent (Life Technologies, USA) followed by DNase digestion using RQ1 RNase free DNase (PROMEGA, USA) according to manufacturer’s instructions. cDNA was synthesized using iScript^TM^ Reverse Transcriptase (BIO-RAD, USA) according to manufacturer’s instructions. Real-time qPCR was performed in 10 µl reactions in an iQ5 instrument (Bio-Rad, CA, USA) using iTaq™ Universal SYBR® Green Supermix (Bio-Rad, CA, USA) and a 400 nM final concentration of each primer from (Supplemental Table 2) Each reaction was run in three technical and three biological replicates. Relative expression levels of genes were assayed using the Pfaffl method (Pfaffl 2001) relative to a previously validated housekeeping reference gene phosphatidylinositol 3- and 4-kinase belonging to the ubiquitin family (PI4K; Medtr3g091400 in MtV4.0, MtrunA17Chr3g0126781 in MtrunA17r5.0-ANR) (Kakar et al., 2008). Data from the three biological replicates were used to estimate the mean and standard error.

### DNA isolation

DNA was isolated from leaf presses made by pressing a leaflet of each plant to a Plant Card (Whatman^TM^, GE Healthcare UK Limited, UK) for long term storage. A 1.2 mm diameter piece of Plant Card was excised and washed with Whatman^TM^ FTA Purification Reagent (GE Healthcare UK Limited, UK) followed by TE-1 buffer according to the manufacturer’s instructions and directly used in PCR. DNA from R108 wild type, used multiple times, was isolated using the DNeasy Plant Mini Kit (Qiagen, Hilden, Germany) as per the manufacturer’s instruction.

### PCR conditions

All amplification except for use in cloning was performed with GoTaq® G2 DNA Polymerase (Promega Corporation, Madison, USA) following manufacturer’s instructions and buffers. Standard PCR conditions used were: 2 min at 95°C, 35 cycles (10 s at 95°C, 15 s at gene specific annealing temperature, 30 s/Kb at 72°C), 5 min at 95°C. When using leaf press as DNA template for amplification, cycles were increased from 35 to 40. The gene specific annealing temperature was 55°C except as noted for primer pairs 3148/3149 (58°C), 3182/3183 (52°C), and 2493/2494 (56°C). Annealing temperatures were predicted using the ‘Tm Calculator’ tool available at https://www.thermofisher.com/. Amplification of inserts for cloning were performed with iProof^TM^ High-Fidelity DNA Polymerase (Bio-Rad Laboratories, USA) and buffers according to the manufacturer’s instructions.

### Imaging

The nodule images in Figure 3 were taken using the Leica Microsystems THUNDER Imager Model Organism with the attached DMC4500 color camera. For visualization of AM fungal structures, colonized roots were cleared in 20% KOH (w/v) at 65°C for two days. Then, they were rinsed three times with ddH2O for at least one day until the roots appeared white, and the pH was neutralized overnight using 1x PBS. Cleared roots were stained in Alexa Fluor® 488 wheat germ agglutinin (WGA)-PBS solution so the final concentration of WGA was 0.2ug/ml. AMF colonization was quantified using the magnified intersections method with a gridded square petri dish (McGonigle et al., 1990). Images were captured with a Leica M205 stereoscope.

### Computational and Statistical analysis

The tree in Figure 1 was generated with (MEGA11 (Tamura et al., 2021) using the maximum likelihood method and tested using Bootstrap for 1000 replicates. The normality of data was tested by fitting a normal curve before performing the statistical tests. Tukey’s comparison for all pairs was used for comparison among more than two groups, and the student’s t-test was used for comparing two groups for normal data. Nonparametric Steel-Dwass comparison for all pairs was used for the analysis was used for non-normal data. Statistical test was performed using JMP Pro 17.1.

**Figure 1:**
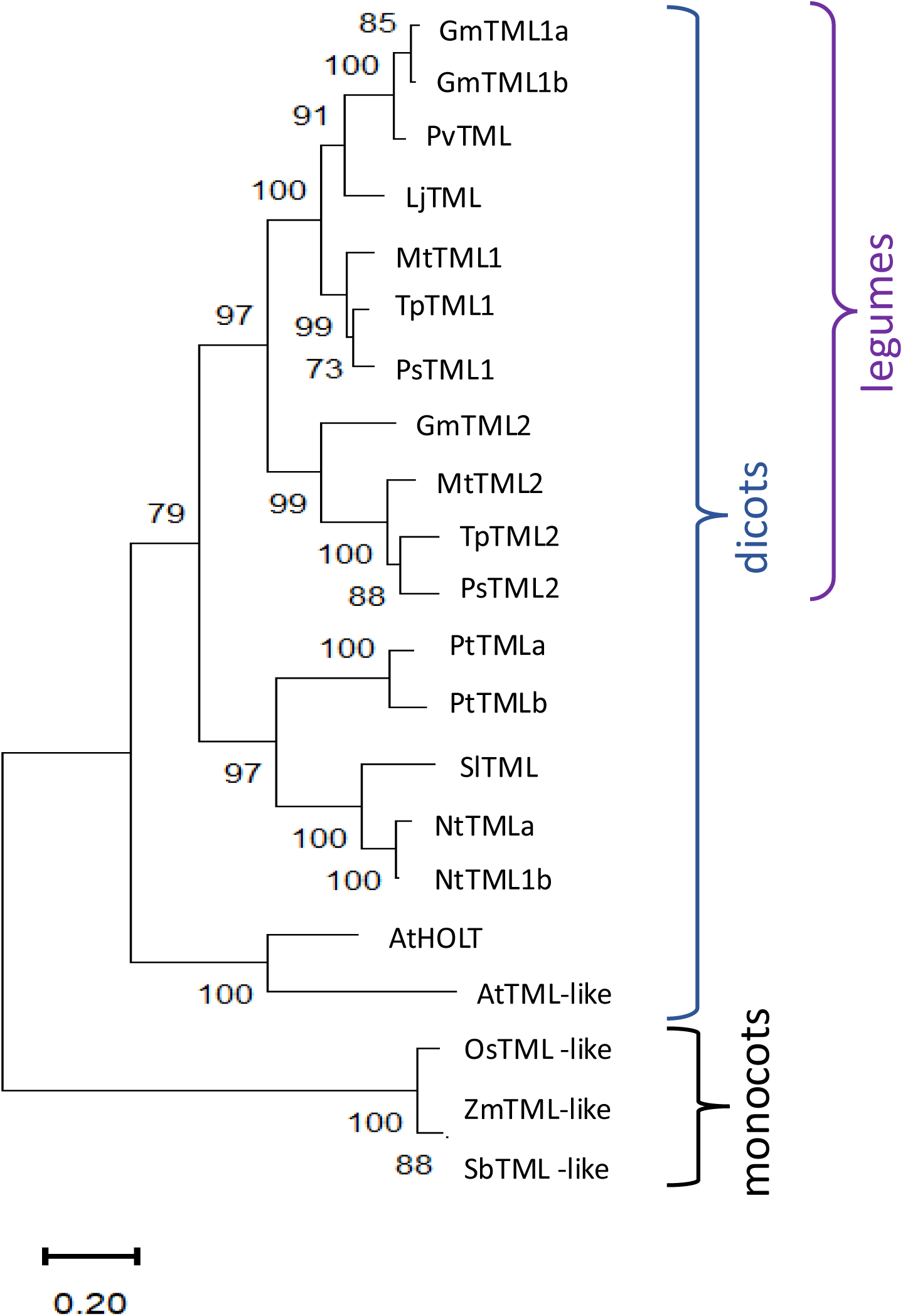
Phylogenetic analysis and structure of TML sequence homologs. Phylogenetic analysis and structure of TML sequence homologs. Analysis by the maximum likelihood method using sequence homologs from public databases (Supplemental Table 1) as described in Methods, in *M. truncatula* (Mt), *Arabidopsis thaliana* (At), *Nicotiana tobacum* (Nt), *Pisum sativum* (Ps), *Trifolium pratense* (Tp), *Glycine max* (Gm), *Phaseolus vulgaris* (Pv), *Solanum lycopersicum* (Sl), *Populus trichocarpa* (Pt), *Zea mays* (Zm), *Oryza sativa* (Os), and *Sorghum bicolor* (Sb). Percentage of replicate trees in which the associated taxa clustered together in the bootstrap test 1000 replicates are shown next to the branches.

## Results

### The number of TML genes in a genome is not correlated with the ability to nodulate or nodule meristem type

Since only one *TML* gene has been identified in *L. japonicus* (Magori et al., 2009) which forms determinate nodules, we wondered if having two copies of *TML*, like *M. truncatula* (Gautrat et al., 2019), was correlated with indeterminate nodulation, and whether multiple copies appear in non-nodulating plants. Figure 1A displays a Maximum Likelihood tree for amino acid sequence homologs of LjTML identified by BLASTP in selected determinate and indeterminate nodule forming legumes, non-legume dicots and monocots. TML proteins from legumes clustered in a separate branch from non-legumes while single TMLs from the monocots rice, maize and sorghum clustered separately forming the tree root; these species have only ∼40% amino acid sequence similarity with LjTML. In cases where non-legumes had two TML proteins, with similar low similarity scores to both TML1 and TML2, the labels are based on previous descriptions or ploidy. Arabidopsis does not form either symbiotic association with rhizobia or mycorrhizae and the Arabidopsis protein *At*HOLT (*HOMOLOG OF LEGUME TML*) has only 51% amino acid sequence identity to LjTML, yet the *AtHOLT* transcript is a target of miR2111 (Hsieh et al., 2009; Pant et al., 2009) and regulates lateral root formation in response to nitrate in the same manner as *Lj*TML. There is no correlation among legumes between *TML* gene number and nodule meristem type. Only one gene encoding a TML protein was identified by BLAST search in the determinate nodulating species *L. japonicus* and *P. vulgaris* (common bean) whereas determinate nodulating soybean has genes encoding three TML proteins. *GmTML1a* and *GmTML1b* are gene duplicates sharing 93.32% sequence identity and the encoded proteins cluster together in the phylogenetic tree, likely the result of the partially diploidized tetraploid soybean genome (Shultz et al., 2006). The same a, b designation used for the soybean duplicate proteins was applied to *N.tabacum,* a recent allotetraploid of *N. sylvestris* and *N. tomentosiformis* (Renny-Byfield et al., 2011) and *P*. *trichocarpa,* which underwent a whole genome duplication approximately 60 million years ago (Dai et al., 2014). The TML2 protein sequences in legumes (*Gm*TML2, *Mt*TML2, *Ps*TML2 and *Tp*TML2) cluster together and branch separately from TML1 sequences, suggesting the two genes encoding TML proteins evolved from a common ancestral duplication in legumes, compared to the independent duplications in other dicots that form clades by species.

### Identification of lines with mutations in MtTML genes

To explore the function of the two TML proteins in *M. truncatula*, we created and isolated mutant lines. The *tml2-1* allele was identified in the NF0679 pool from the *Tnt1* insertional mutagenesis library and isolated as a homozygous line (see Methods). Additional mutant alleles of *MtTML1* and *MtTML2,* as well as plants containing mutations in both genes, were generated by CRISPR-Cas9 mutagenesis of embryonic callus tissue followed by regeneration of whole plants in the R108 ecotype (see Methods). Multiple combinations of heterozygous and biallelic mutations were identified and two individual alleles of each gene, as well as two double mutant combinations were used in this study (Figure 2; Supplemental Figure 1). The *tml1* mutants created by CRISPR include a deletion in *tml1-1* resulting a premature stop codon, which was recovered in the creation of a double mutant and only characterized in the *tml1-1 tml2-2* double mutant. The two other alleles of *tml1* were isolated and characterized as single mutants: *tml1-2*, which removes 111 aa from the N terminus of the protein, and *tml1-4*, a deletion eliminating the start codon. The *tml2* single mutants include a CRISPR-generated deletion and the above-described *Tnt1* insertion, both resulting in stops early in the protein. As a result, the *tml1-1 tml2-2* double mutant is the combination of two early stops and the *tml1-2 tml2-2* double mutant is the combination of an early deletion and a stop. All stops occur prior to or early in the F-box coding region, so that any resulting truncated protein is not expected to contain the F-box, NLS, and kelch repeats (Figure 2). The binding sites for miR2111 in *MtTML1* and *MtTML2* start at 143bp and 134bp into the CDS respectively, and the binding site is lost only in the *tml1-4* mutant. (Supplemental Figure 1).

**Figure 2.**
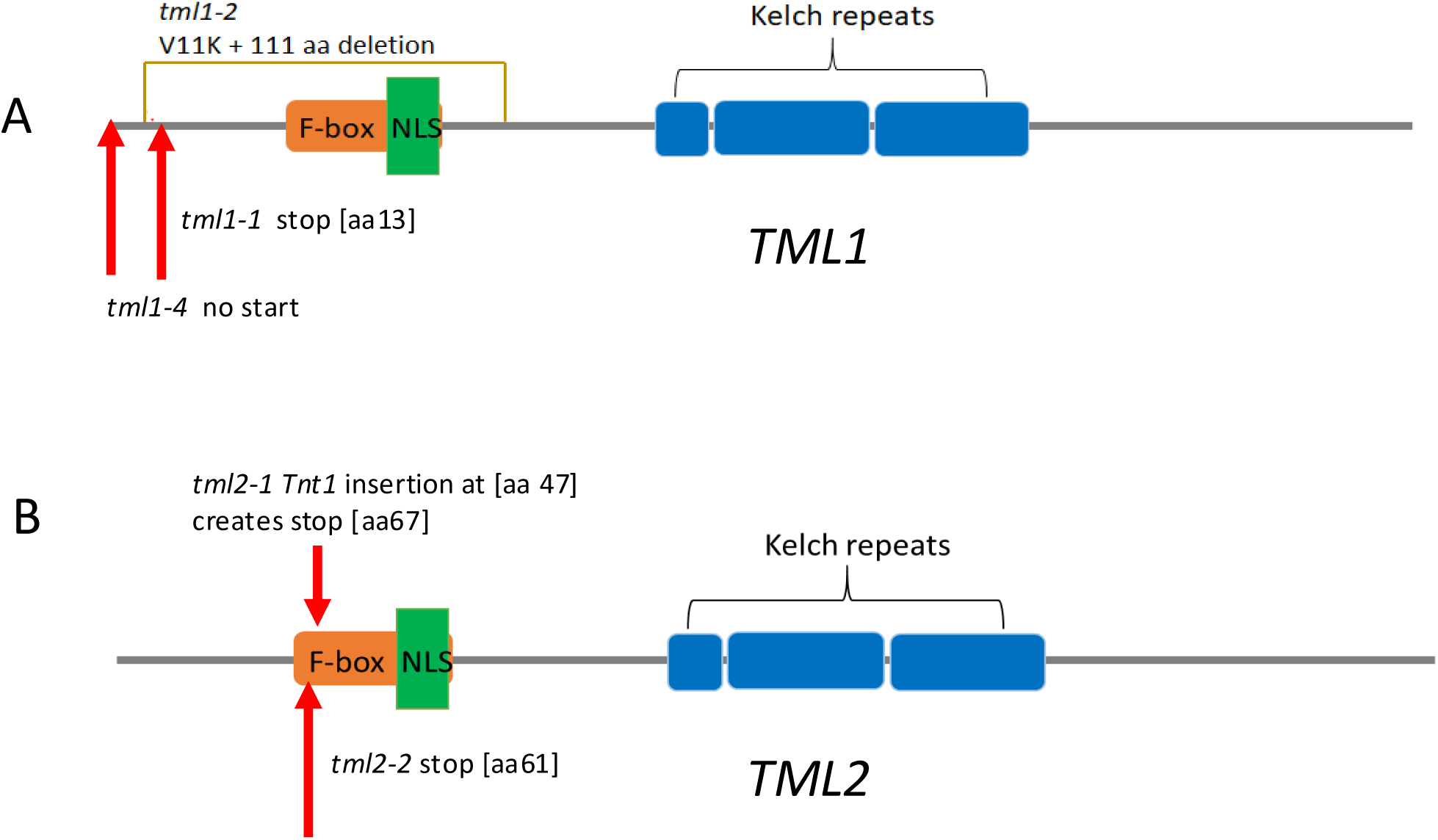
*TML* alleles used in this work. Diagrams are based on the translation of the R108 genotype sequence of Medtr7g029290 (*MtTML1*) and Medtr6g023805 (*MtTML2*). Protein features are shown by colored boxes: F-box in orange, kelch repeats in blue, Nuclear Localization signal (NLS) in green. miRNA2111 binding sites are in the coding sequence (CDS) indicated in Supplemental Figure 1. (A) *TML1* alleles. *tml1-2* is a 334 bp deletion with the addition of an A that results in a V to K change at amino acid 11, and deletion of 111 amino acids, removing the F-box and NLS. *tml1-4* is a 104 bp deletion in the 5’ UTR and beginning of the coding sequence that removes the start codon and the first 20 amino acids. (B) *tml2-1* is a *Tnt1* insertion 119 bp from the start of the coding region resulting in addition of 20 new amino acids starting at position 47 before eventually terminating the protein at position 67. *tml2-2* is an insertion of a C, creating a stop codon at amino acid position 60 in the F box.

### MtTML1 and MtTML2 single mutants have slightly increased nodule numbers while double mutants hypernodulate

The nodule number phenotypes of plants carrying mutant *tml1, tml2*, and *tml1 tml2* double mutant alleles were determined 10 days post inoculation (dpi) with *Sinorhizobium meliloti* RM41 in an aeroponic system and compared to wild type R108 plants and hypernodulating *sunn-5* plants (described in (Crook et al., 2016; Nowak et al., 2019)). Plants carrying a mutation in a single *MtTML* gene had more nodules than wild type plants, while plants with mutations in both *MtTML* genes had many more nodules and shorter roots than wild type plants (Figure 3A). All single mutant lines formed pink nodules similar in size to wild-type plants (Figure 3B-F) whereas the double mutant plants formed small, fused nodules in a wider area of the root (Figure 3G-H). Most nodules in the double mutants were white at 10 dpi, except for a few slightly pink nodules indicating leghemoglobin production. No obvious shoot phenotype was observed in any line.

**Figure 3.**
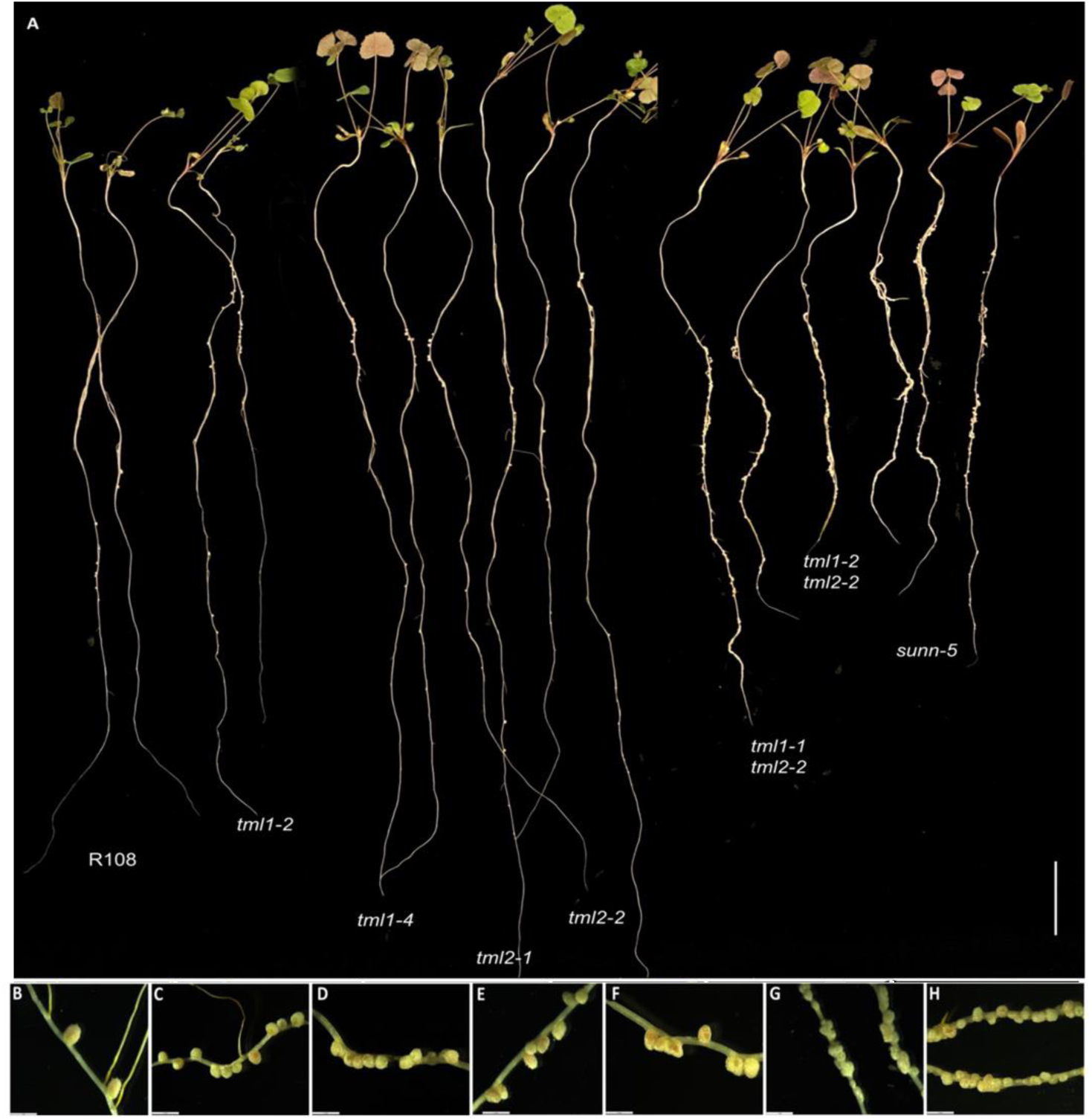
Phenotype of single and double mutant alleles in *MtTML1* and *MtTML2* genes. (A) whole plants displaying root nodules at 10dpi in an aeroponic system inoculated with *Sinorhizobium meliloti* strain RM41. Two representative plants per genotype L to R: R108 (wild type), *tml1-2*, *tml1-4*, *tml2-1*, *tml2-2*, *tml1-1 tml2-2*, *tml1-2 tml2-2* and *sunn-5.* Scale bar = 2 cm. (B-H) magnification of nodules from a representative plant for genotypes (B) wild type R108, (C) *tml1-2*, (D) *tml1-4*, (E) *tml2-1*, (F) *tml2-2*, (G) *tml1-1 tml2-2* and (H) *tml1-2 tml2-2.* A single root was folded in (H) to show wider area of nodulation in same picture; B to H scale bar = 2mm.

All mutants in a single *MtTML* gene formed a significantly higher number of nodules than the wild type but less than the known hypernodulating *sunn-5* mutant (p ≤ 0.01, Steel-Dwass Method of pairwise comparison; Figure 4A). The evaluation of nodules per cm of root length followed the same pattern as the total number of nodules (Figure 4B). However, the double mutants formed a significantly higher number of total nodules and nodules per cm of root length than both the wild type and *sunn-5* mutant (Figure 4 C and D). The single mutants on average formed 2-3-fold more nodules while the double mutants formed greater than 20-fold more nodules compared to the wildtype (Figure 4).

**Figure 4.**
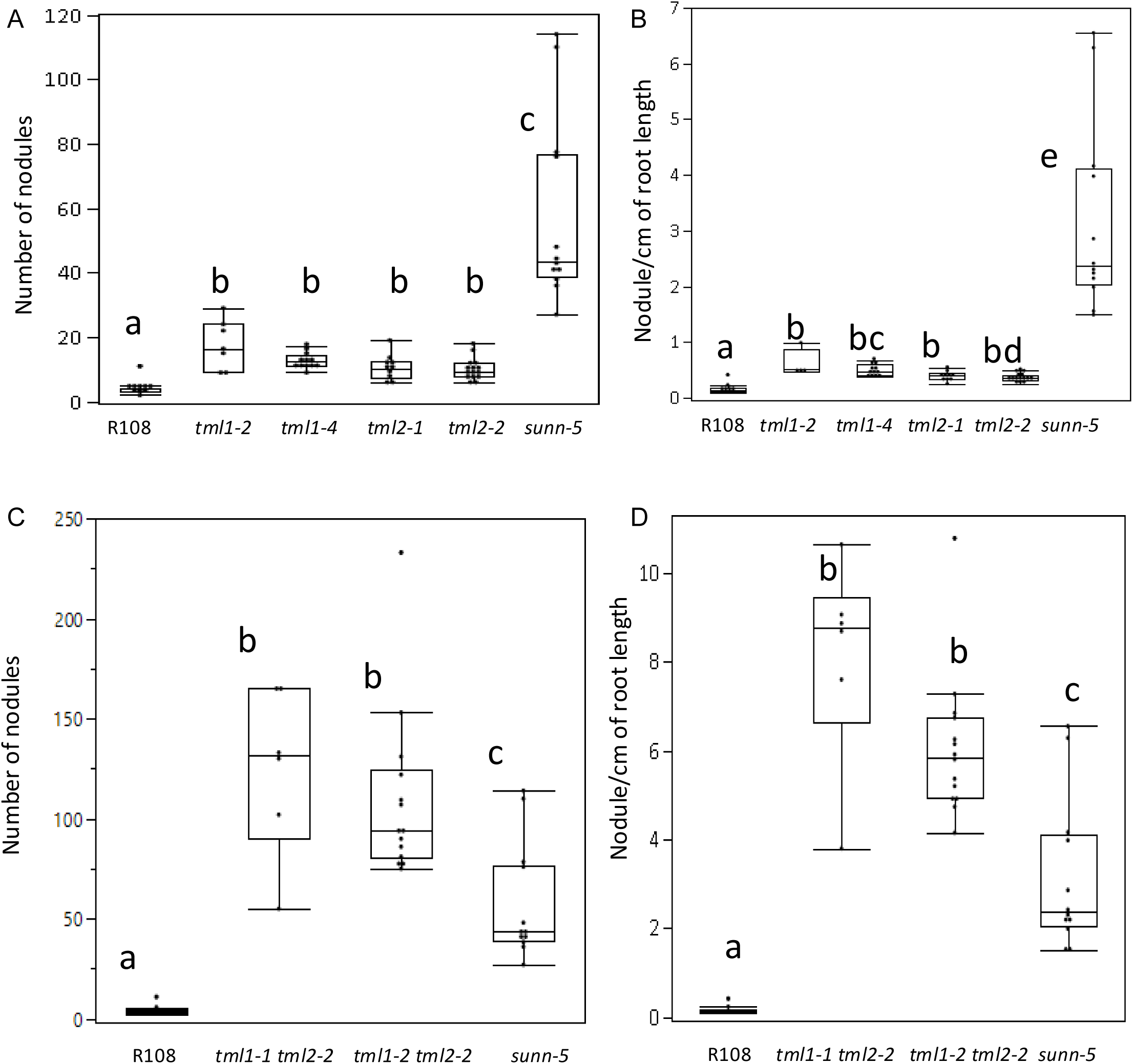
Nodulation phenotypes of single and double mutant lines in *MtTML1* and *MtTML2* genes. (A) Nodule number and (B) nodules per cm root length at 10dpi of wild type, single mutant, *sunn-5* plants from the same experiment in Figure 3. N=12 plants for R108 wild type and N=15 plants for *sunn-5* in all graphs. N =7 plants for *tml1-2*, N = 10 for *tml1-4,* N=11 for *tml2-1, and* N=19 for *tml2-2*. Groups not connected by the same letter are statistically different. (C) Nodule number and (D) nodules per cm root length at 10dpi of wild type, double mutant and *sunn-5* plants from the same experiment in Figure 3. Groups not connected by the same letter are statistically different. N=8 plants for *tml1-1 tml 2-2* and N=14 for the *tml1-2 tml2-2* double mutant. Significant difference between means were tested Tukey’s comparison for all pairs or Nonparametric Steel-Dwass comparison (see Methods for detailed description). Dots indicate values for individual plants.

### MtTMLs differ in their spatiotemporal expression during nodule development

Using the ePlant resource for early nodulation (Schnabel et al. 2023b) and past transcriptomics work in our lab (Schnabel et al., 2023) we examined the expression of the two *MtTML* genes in root segments during the first 72 hours responding to rhizobia (Figure 5). The genes are expressed in wild-type plants at low levels at the 0 hr timepoint, which is 4-day old N starved plants in our experimental system, but expression rises in the vascular tissue for both genes at 12 and 24 hpi (Figure 5A). By 48 hpi when the nodules are beginning to form, *MtTML1* is expressed at a similar level in the vascular, cortical, and epidermal laser captured cells as well as the developing nodule cells, while *MtTML2* increases to a higher level in the vascular and inner cortical cells at the xylem poles and does not increase in the nodule. By the time the nodule has organized and begun to emerge at 72 hpi, this difference in pattern is reinforced, with *MtTML1* highly expressed in the nodule and *MtTML2* expressed in the nodule at the same low level as in tissues of uninoculated plants. Because of the way the laser capture data is collected and displayed, color indicates expression somewhere within the cell types, rather than throughout the area colorized. Examined at the level of single cell RNASeq over a time course of combined experiments harvested at time points at 0, 48 and 96 hpi, the combined transcriptomics data of individual cells responding to rhizobia showed both *MtTML* genes are expressed in the pericycle cells and vascular bundles, but *MtTML1* is expressed in different cell lineages than *MtTML2* in response to rhizobia (Pereira et al., 2024). Observation of expression levels in undissected root segments over time revealed that both *MtTML* genes are induced 2-3 fold in wildtype responding to rhizobia over 72 hours, as well as in the hypernodulation mutant *sunn-4* (Schnabel et al., 2023). However, *MtTML1* expression does not begin to decrease over this time frame in the *sunn-4* mutant, while *MtTML2* does. Taken together, the data imply that *MtTML1* and *MtTML2* have a spatiotemporal effect on nodulation; *MtTML1* has higher expression in cortical cells and nodules, whereas *MtTML2* is expressed mainly in the pericycle and vasculature.

**Figure 5.**
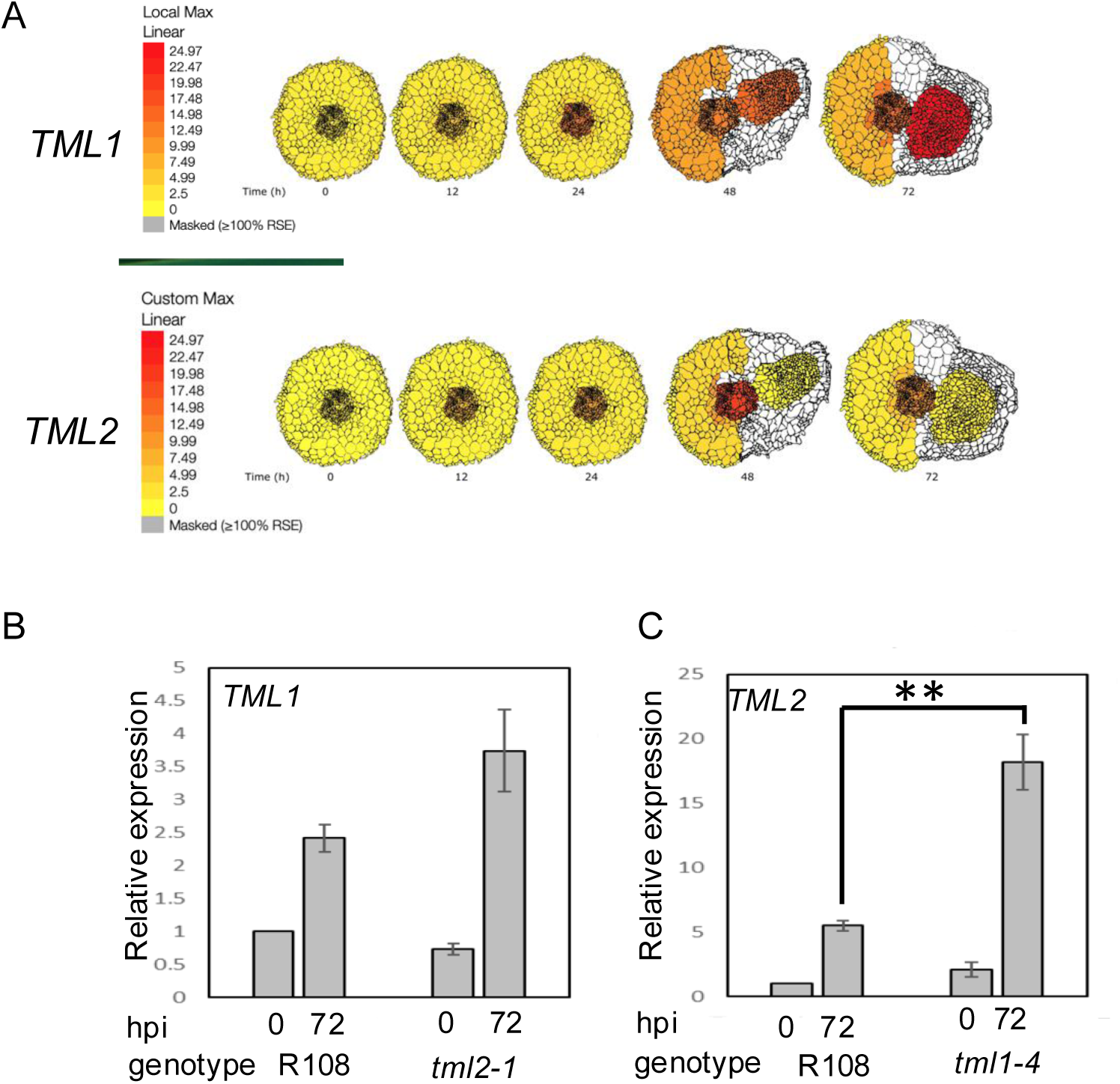
Spatiotemporal and differential expression of *MtTML* genes over 72 hours in plants responding to rhizobia. (A) Diagrams produced from ePlant resource (Schnabel et al., 2023b) with gradient display adjusted to identical color representation to allow comparison of the expression levels as well as localization of *TML1* and *TML2* across time in wild type roots responding to rhizobia up to 72 hpi. (B) Relative expression of *TML1* in a *tml2-2* mutant at 0 and 72 hours post inoculation (hpi), normalized to wild type at 0 hpi. (C) Relative expression of *TML2* in a *tml1-4* mutant at 0 and 72 hours post inoculation normalized to wild type at 0 hpi. Note Y axis in B is 5 times the Y axis in A. ** p< 0.05. Each graph is the results from 3 technical replicates each of 3 biological replicates from 10 plants each, compared to the reference gene PIK (Kakar et al., 2008).

### tml1 mutants show evidence of transcriptional adaptation

Genetic compensation can occur by several mechanisms but transcriptional adaptation, in which degradation of the mutant mRNA transcript triggers increased expression of a paralog or family member (El-Brolosy et al., 2019), is straightforward to test. Using Real Time Quantitative PCR, we measured expression of one wildtype *MtTML* gene when the other *MtTML* was mutated. In the roots of wild-type R108 and plants containing either a *tml2* or *tml1* mutant transcript, there is no statistical difference in expression of the wild-type *MtTML* message in the absence of rhizobia (0 hpi), and all plants showed statistical increases in expression of the examined transcript from the uninoculated level (0 hpi) compared to 72h after inoculation with rhizobia (72 hpi) (p<0.05) (Figure 5 B, C). In the *tml2* mutant, the expression level of *MtTML1* at 72 hpi is statistically equivalent to wildtype at 72 hpi, showing no evidence of genetic compensation (Figure 5B). However, in the *tml1* mutant, the expression level of *MtTML2* is increased from that of wild type at 72 hpi (p<0.05) (Figure 5C), suggesting that the loss of a functional copy of *MtTML1* increases the expression of *MtTML2*, but not vice versa. In wild-type plants, the fold changes of *MtTML1* at 72 hpi are half that of *MtTML2*, which could make differences in *MtTML1* expression significance harder to detect.

### Phenotypic effects of heterozygosity appear in segregation of double TML mutants

Following the observation that expression of the wild-type *MtTML2* gene increased in a plant containing a mutation in *MtTML1* but did not rescue the nodule number phenotype of the *tml1* mutant, we wondered if there is effect on the nodulation due to heterozygosity of mutant allele. While an intermediate nodule number for plants heterozygous for a *tml* mutation might be difficult to detect given the small size of the effect on nodule number of a single mutation (2-3-fold change increase in single mutants; Figure 4A), the magnitude of the phenotype of double mutants could allow a nodule number difference to be detectable when plants are homozygous for one mutant allele and heterozygous for the other.

We examined two populations of progeny obtained from selfing plants homozygous for one *tml* mutation and heterozygous for the other, *tml1-2*/TML1 *tml2-2* and *tml1-2 tml2-2*/TML2. A discrete recessive phenotype should segregate nodule number phenotypes in a 3:1 Mendelian ratio. Instead, both populations segregated a broad distribution of nodule number phenotypes (Figure 6). To test our hypothesis that this distribution of phenotypes is due to changes in the inheritance of the segregating wild type copy, we used the progeny of plants homozygous for *tml2-2* and segregating *tml1-2*. Out of 141 progenies, a random set of 84 (∼60%) of the population was tested by PCR. The PCR tested progeny were verified for the expected 1:2:1 genotypic segregation using Pearson’s Chi square test (χ^2^ = 7.13, with 2 degrees of freedom). The analysis revealed plants wild type for the segregating allele had a mean of 22.5 nodules, plants heterozygous for the segregating allele had a mean of 43.8 nodules, and plants homozygous for both mutant alleles had a mean of 100.3 nodules. Each of the three groups were significantly different from each other in Tukey’s all pair comparison (p<0.001) (Supplemental Figure 2), supporting a phenotypic effect of heterozygosity on the nodule number phenotype.

**Figure 6.**
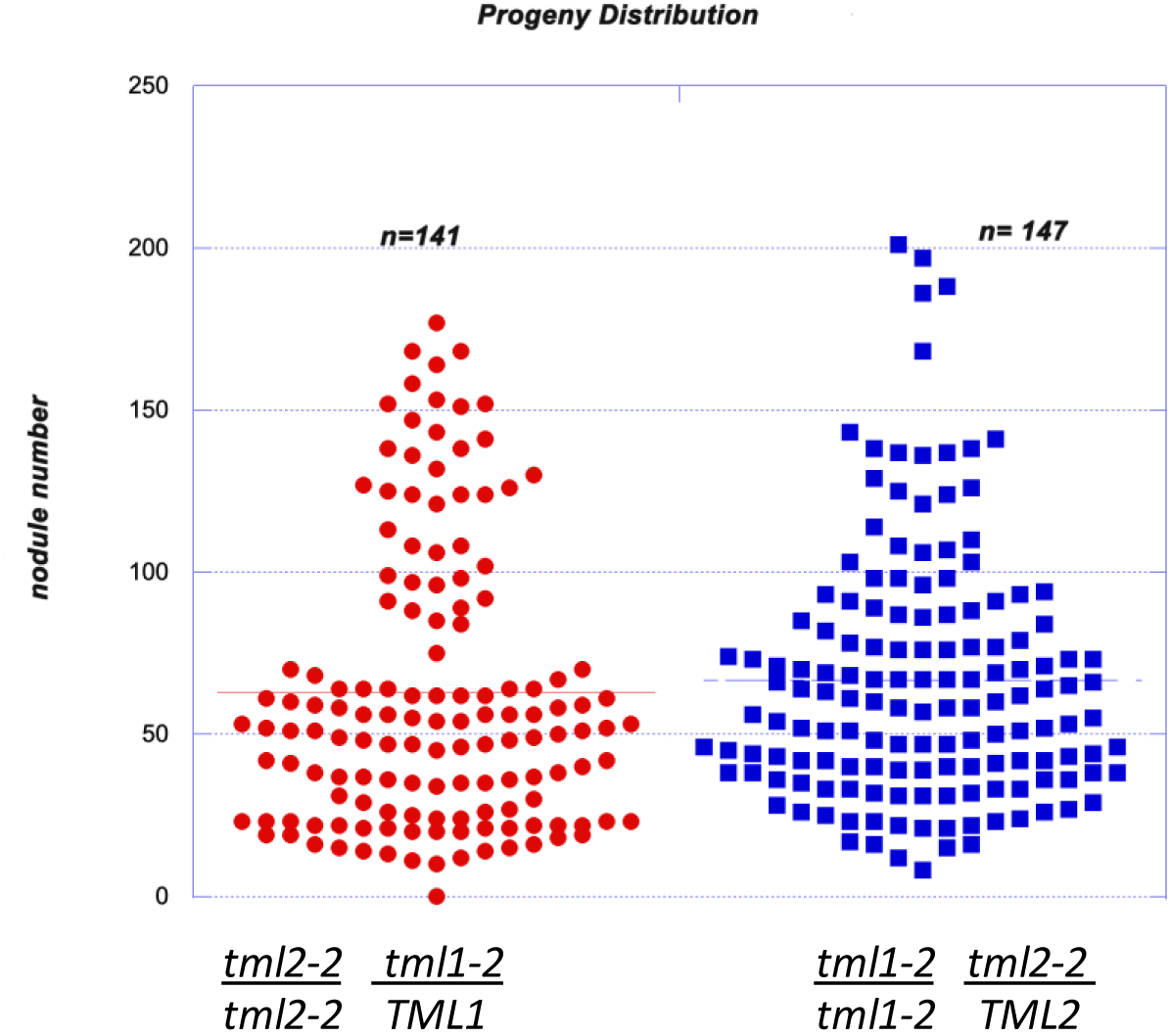
Distribution of nodulation number phenotypes suggests heterozygous effect. Distribution plots of two populations of progeny from selfing of plant on X axis. Red dots are individual plant nodule numbers from *tml2-2* mutant plants segregating one wild type *TML1* allele, blue squares are *tml1-2* mutants segregating one wild type *TML2* allele.

### tml double mutants have minor effects on arbuscular mycorrhizal symbiosis

Many AON mutants also have hyper-mycorrhizal phenotypes, and *miRNA2111* is induced upon phosphate starvation even in Arabidopsis (Hsieh et al., 2009), which does not form symbiotic associations with rhizobia or AM fungi. Expecting an increase in AM fungal root colonization, we tested the *tml1-1, tml2-2* and *tml1-2, tml2-2* double mutants for a mycorrhizal phenotype with the AM fungus *Rhizophagus irregularis*. No overall root length colonization difference between double mutants and wildtypes was observed under our growth conditions at 4.5 weeks post inoculation (Figure 7A). However, in the same experiment at 6 weeks post inoculation, a small but statistically significant increase in AM fungal root length colonization was observed for the *tml1-2 tml2-2* double mutant relative to R108 wild-type controls, but not the *tml1-1 2-2* allele (Figure 7A). Arbuscule morphology appeared normal in the mutants (Fig. 7 B-D). Together, this data suggests that *MtTML1* and *MtTML2* might play a minor role in AM symbiosis although their impact appears to depend on the timepoint measured and the mutant allele.

**Figure 7.**
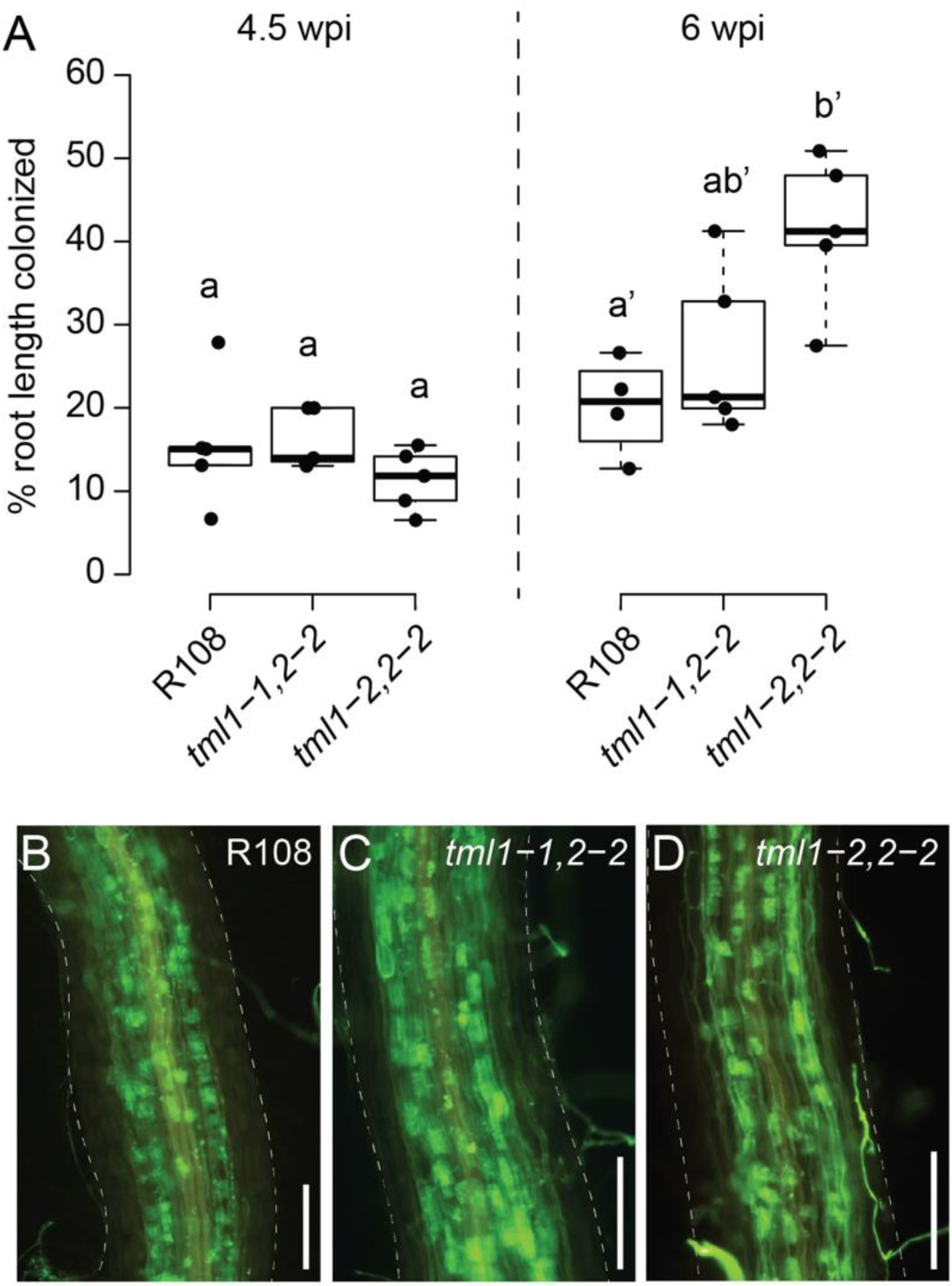
Arbuscular mycorrhiza phenotype of *tml1, tml2* double mutants. A) Overall root length colonization in R108, *tml1-1, tml2-2,* and *tml1-1, tml2-2* is similar at 4.5 weeks post inoculation (wpi), whereas at 6 wpi increased colonization levels were observed in the *tml1-2, tml2-2* double mutants relative to R108 controls. Statistical differences were calculated separately for each timepoint (ANOVA followed by Tukey’s HSD; different letters denote significant differences in pairwise comparisons with p<0.05). B) Representative image of *Rhizophagus irregularis* symbiotic structures in an R108 wild-type root (6wpi). C) Representative image of a *R. irregularis*-colonized root of a *tml1-1, tml 2-2* double mutant (6wpi). D) Representative image of a *R. irregularis*-colonized root of a *tml1-2, tml2-2* double mutant (6wpi). B-D) *R. irregularis* fungal structures are visualized with WGA-Alexafluor488. *M. truncatula* root is outlined with a dashed line. Scale bar: 250μm.

## Discussion

While the *TML* gene was first identified and investigated in *L. japonicus* (Ishikawa et al., 2008; Yokota et al., 2009, (Magori et al., 2009), legumes and most dicots examined contain two copies of a *TML*-like gene, with tomato and *L. japonicus* as outliers (Takahara et al., 2013). Based on our ability to only identify one *TML* copy in the monocots we analyzed, we suspect an ancestral gene duplication early in the legume lineage, followed by the loss of a copy in *L. japonicus*. Dicots containing more than one copy appear to be the result of genome duplications and are more closely related in sequence to each other than to either legume *TML*, hence the a, b labeling in Figure 1. As is common for many genes in soybean, the *G. max* genome contains two copies of *TML1*. Since monocots and dicots diverged approximately 200 million years ago (Wolfe et al., 1989), and legumes even later, enough time has passed for the genes to have adopted divergent functions, but this work suggests any divergence within legumes is complex, and the genes have evolved to give synergistic effects in some of the symbiotic phenotypes tested and none in others.

Given the moderate effect of RNAi against the *M. truncatula TMLs* on nodulation phenotypes (Gautrat et al., 2019), we expected the strong genetic mutants created in this study (stop codons early in the gene) to have a larger effect on phenotype than observed. However, all single mutations, predicted to result in highly truncated proteins if translated, had small effects on nodule number, displaying only 2-3 times the number of nodules as wild type plants (Figure 4). In sharp contrast, mutation in the single copy of *TML* in *L. japonicus* results in eight times more nodules than wild type (Magori et al., 2009). When both copies of *TML* were disrupted in *M. truncatula,* the nodule number phenotype was logarithmically increased, exceeding that observed for *sunn* mutants (Figure 4) and for the *L. japonicus tml* mutant (Magori et al., 2009). In our aeroponics system (Cai et al., 2023), all plants encounter rhizobia at the same time, leading to a distinct zone of nodulation even in AON mutants such as *sunn* and *rdn1* (Schnabel et al., 2005; Schnabel et al., 2011). Interestingly, while we observed the same effect in single *tml* mutants, the double mutants nodulated along the entire root (Figure 3) suggesting that competence to nodulate may be more than just on/off switch (Laffont and Frugier, 2023) and other characteristics such as the extent of the nodulation zone or the speed of nodule development could be regulated by a signal dependent on both TML proteins.

The observation of transcriptional adaptation in *tml1* mutants (increased expression of the *MtTML2* gene during nodulation (Figure 5C)) could suggest some overlap in function between the two genes. *MtTML1* is expressed in the nodules at 72 hpi (likely the nodule vasculature based on (Pereira et al., 2024)) and *MtTML2* is not (Figure 5A). Yet, the loss of *MtTML1* results in increased expression of *MtTML2*, which is normally not expressed in the nodules (Figure 5C). However, identification of a slightly increased nodule number phenotype in *tml* single mutants (Figure 4) and *TML* heterozygotes (Supplemental Figure 2) also suggests that the two genes are not completely redundant. As noted in the description of the results for Figure 5, because the fold change observed in *MtTML1* expression at 72 hpi is much smaller than that of *MtTML2,* genetic compensation for *MtTML1* in *tml2* mutants could exist but be too small for a significant change in expression to be observed in our experiment. However, even if genetic compensation does not occur, an overlap of function could explain the small effect on nodule number in single mutants. Additionally, the differences in mean nodule number for *tml1-2 TML1* heterozygotes segregating in a *tml2-2* background versus plants carrying a wild type allele in the same background suggest an effect of the amount of wild-type *TML* message on the number of nodules formed. Combined with the additional observation of a similar distribution of nodule number in *tml2-2 TML2* heterozygotes segregating in a *tml1-2* background (Figure 6, Supplemental Figure 2), the data indicate that the combined level of wild-type *MtTML* message (*MtTML1* + *MtTML2*) may affect the number of nodules formed.

Synergistic gene interaction can have multiple causes (Pérez-Pérez et al., 2009). The single and double mutant alleles in this analysis contain premature stop codons (Figure 2), predicted to result in a truncated protein without any functional domains. The prediction of complete loss of function (null) mutations thus rules out that the synergistic phenotype is the result of hypomorphic alleles. Synergy can also be the result of mutations in in two unrelated genes acting in parallel pathway that converge at a node, or completely or partially redundant paralogous genes functioning in the same step of a linear pathway or mutation. The *MtTML* genes are not unrelated, ruling out the first cause. The second cause is ruled out also- *MtTML1* and *MtTML2* genes cannot be completely redundant genes acting in same spatiotemporal step because mutation in one *MtTML* gene is not completely complemented by the presence of a wild-type copy of the other *MtTML* gene. However, partial redundancy of two genes functioning in the same step can be a result of a differential expression pattern in which genes perform same function in different cellular locations or in different tissue types. This is more likely, as tissue expression data from wildtype (Figure 5A) shows an increase of both *MtTML1* and *MtTML2* transcripts in response to rhizobia over 72 hpi, but there are small differences in both tissues and levels of expression of the *MtTML* genes at 72 hpi. Both *MtTML1* and *MtTML2* are detected at 24 hpi in the vasculature (pericycle), however *MtTML1* is expressed in the developing nodules and cortex at 48 and 72 hpi as well, while *MtTML2* expression remains vascular. The result is different transcript levels in different tissues, suggesting a tissue-specific role at least for *MtTML1*. Since the root segment expression pattern (Schnabel et al., 2023) also shows a difference in temporal expression pattern between *MtTML1* and *MtTML2*, this could indicate similar functions but a different spatial role for *MtTML1*.

Finally, synergy can be observed when the products of two genes interact in a multimeric protein complex; mutation of one component leads to fewer monomers resulting in fewer functional protein complex in a dose-dependent manner. While purely speculative, one possible explanation for both the individual phenotypes and the synergy observed is that the two proteins form homo-or heterodimers as part of their downstream signaling. It is also possible the two proteins might function together bound to a third target, such as a promoter, a transcription factor, or to provide the localization signal to bring a factor into the nucleus.

Given that TML regulation of AON is predicted to result from destruction of the mRNA by miR2111, we suggest that the synergy comes from a dosage dependence of the wild-type message combined with genetic compensation. The lower transcript levels of both *MtTML1* and *MtTML2* in roots under low N conditions is correlated with root competence to nodulation (0 hpi in Figure 5) whereas higher transcript levels of both genes is correlated with inhibition of nodulation after rhizobial inoculation later in time (Gautrat et al., 2020).

A key factor in the AON model is miR2111 produced in the shoot and transported to the root (Okuma and Kawaguchi, 2021; Bashyal et al., 2023). Multiple genes encode miR2111, which is predicted to bind to *MtTML* RNA and decrease RNA levels versus preventing transcription. (Gautrat et al., 2020). When N is abundant, *MtCLE35* is induced in the roots and represses miR2111 in the shoots and roots, resulting in low abundance of miR2111 in the roots and thus higher levels of *MtTML* message (Tsikou et al., 2018; Gautrat et al., 2020; Moreau et al., 2021). Low N is signaled through systemic CEP7/CRA2 peptide/receptor signaling and reduction of systemic MtCLE35/SUNN peptide/receptor signaling. Both signals are transduced through miR2111 levels dropping and *MtTML* message levels rising, signaling competence to nodulate (Moreau et al., 2021; Laffont and Frugier, 2023). When competent plants begin to nodulate, *MtTML* levels continue to rise as nodulation begins along with MtCLE13/SUNN signaling (Schnabel et al., 2023) and fall as nodulation proceeds and miR2111 levels rise, a result of the MtCLE13/SUNN signal AON feedback.

The current AON model is highly simplified, involving signal transduction in both space and time based on evidence gathered by multiple groups in multiple systems and species. We do not know the half-life of the MtTML proteins, which would affect the length of time between detecting decreased mRNA and the action of the MtTML proteins in halting nodulation; split root analysis suggests the AON systemic signal has an effect on nodule number within 72 hpi in *M. truncatula* (Kassaw et al., 2015) but no decreased *TML* expression is detected during this time in our system. What appear to be differences in *MtTML* expression results between our work and others can explained by differences in timing and set up between experiments - the decreased *MtTML* expression data in (Gautrat et al., 2020) was collected at 5 dpi in whole roots on plates versus the increase of *MtTML1* in (Schnabel et al., 2023) at 3 dpi in nodulating segments of roots in an aeroponic chamber. Our results in general support the position of TML in the AON model, but also provide evidence that the functions of the individual *MtTML* genes are not redundant.

AON and AOM share multiple common components, including common LRR-RLKs that perceive distinct CLE peptides specific to nodulation and AM symbiosis, respectively (Roy and Müller, 2022). When legumes are inoculated with rhizobia and AM fungi, the two symbioses influence each other (Catford et al., 2003), potentially by competing for carbon. However, we previously reported that the AON regulator MtCLE13 does not influence AM symbiosis, suggesting AON and AOM have symbiosis-specific signaling outcomes (Müller et al., 2019). Because the distinct CLE signals mediating AON, AOM, and P and N regulation of nodulation and AM symbiosis, converge at the same LRR-RLK SUNN but result in at least partially distinct outcomes (Figure 8), it appears plants can distinguish between the signals. Along these lines, the *M. truncatula* pseudokinase *MtCORYNE* was found to be involved in mediating AON signaling downstream of *MtCLE12* and *MtCLE13* (Nowak et al., 2019), whereas it is largely dispensable for AOM signaling by *MtCLE53* (Orosz et al., 2024). Similarly, while our results demonstrate that *M. truncatula TML1* and *TML2* are important components of AON signaling, they seem to play a weaker role in AOM. We only observed small increases in AM fungal root colonization in the *tml1-2, 2-2* double mutant (in which *MtTML1* is disrupted by a 111 bp deletion) and no detectable increase when using the *tml1-1, 2-2* allele, which is caused by a nonsense mutation near the N-terminus of the protein (Fig. 2, Fig. 7), in which case occasional stop codon read-through may produce a truncated but at least partially functional MtTML1 protein (Zhang et al., 2024). Such phenotypic difference between the two alleles was not observed in our nodulation experiments (Fig. 4), presumably because the nodulation phenotype is much stronger than the relatively mild AM symbiosis phenotype.

AON and AOM share multiple common components, including common LRR-RLKs that perceive distinct CLE peptides specific to nodulation and AM symbiosis, respectively (Roy and Müller, 2022). In addition, P and N regulate nodulation and AM symbiosis through similar pathways converging at the same LRR-RLK SUNN (Roy and Müller, 2022). Although we found only a weak phenotype in one of the double mutant alleles tested, our results suggest *MtTML* genes play at least a small role in AM symbiosis regulation. One tempting hypothesis is that the effect of TML on AM symbiosis stems through intersection with nutrient homeostasis signaling. miR2111, which targets *TML*, is also regulated by N- and P-starvation-induced CEP peptides that signal through the RLK CRA2 to regulate nodulation and AM symbiosis, respectively (Laffont et al., 2019; Laffont et al., 2020; Ivanovici, 2021; Ivanovici et al., 2023; Lepetit and Brouquisse, 2023).

This study uncovered the non-redundant role of MtTML1 and MtTML2 in AON and AOM and opens a new avenue for definitive additional experiments in *M. truncatula* to refine and fill in gaps in the model with the *MtTML* mutants we have created. It is unknown how *SUNN* signaling results in changes in miR2111 expression in the shoot or whether the destruction of *TML* RNA by miR2111 binding in the root affects further *TML* expression. It is unknown how the TML proteins exert their effects and through what other proteins (though the presence of nuclear localization signals might suggest interaction with transcription factors). It is possible there are regulatory steps that involve loading and unloading in the vasculature or import to the nucleus that involve other proteins TML reacts with. The mutants described here will allow these and other important questions to be investigated.

## Acknowledgement & Funding

We acknowledge Clemson Light Imaging Facility and Dr. Terri Bruce, Rhonda Powell and J. Conrad Epps for training and assistance with the Leica THUNDER Imager equipment use. We acknowledge Hemani Patel, an undergraduate mentored by Diptee Chaulagain, for the PCR work on segregating single mutants. We thank Dr. Li Wen for teaching *M. truncatula* crossing to DC and Dr. Yuanling Chen for tissue culture tips and troubleshooting. This work was supported by NSF 1733470 to JF and USDA-NIFA 2022−67013−36881 to LMM.

## Authorship contributions

D.C. designed the research, performed research, analyzed data

E.S. designed the research, performed research

E.X.L. performed research

M.K. performed research

L.M.M. designed the research, performed research; analyzed data

J.F. designed the research, performed research; analyzed data

All authors contributed to drafting the work or revising it critically for important intellectual content.

## Supplemental Materials

**Supplemental Figure 1. Multiplexed targeted CRISPR/Cas9 mutagenesis in *TML* genes using Csy4 plasmid**. (A) Schematic diagram of polycistronic gene construct used in pDIRECT_23C+TML1/2 for making tml1/tml2 double mutants and (B) in pDIRECT_23C+TML1 for making tml1 mutant (C) Maps of *MtTML1* and *MtTML2* genes showing the location of targets. PAM for each target is underlined. Blue boxes indicate CDS, grey boxes indicate UTR, red rectangles indicate predicted miRNA2111 binding sites, and lines indicate introns; arrow at the right end of grey box points to 3’ end of gene. (D) CRISPR effects on DNA resulting in *tml1* alleles, displayed in relation to the target sequences (yellow) and PAM sequences (green). Base changes and deletions are in red. The *tml1-4* allele has a 107 bp deletion that removes 48bp upstream of the transcription start site and 59bp in the CDS, including target 1. (E) CRISPR effects on DNA in the *tml2* allele, displayed in relation to the target sequences (yellow) and PAM sequences (green). Base changes and deletions are in red.

**Supplemental Figure 2. Molecular genotype analysis and photographs supporting heterozygous effect.** (A) Testing 60% (84) of the total population of 141 (indicated in red dots in panel 1) broken down in panels by molecular genotype determined by PCR. Blue dots (panel 2) are homozygous wild type *TML1,* green dots (panel 3) are heterozygous *tml1/TML1* and black dots (panel 4) are homozygous *tml1/tml1*. Lines indicate median nodule number for each group. (B) Box plot showing the distribution of nodule number in each genotype. Groups indicated by different letters are significantly different as tested by Tukey all pair test (p<0.001). (C) One representative plant per genotype L to R: R108 (wild type), *tml1-2*, *tml1-2 tml2-2* het, *tml1-2 tml2-2* and (D) R108 (wild type), *tml2-2*, *tml1-2* het *tml2-2*, and *tml1-2 tml2-2*. Scale bar = 2 cm.

**Supplemental Table 1.** Genes used in Maximum Likelihood tree in Figure 1

**Supplemental Table 2.** Primers used in this manuscript

