## Supplemental Figure 1 for "*TML1* AND *TML2* SYNERGISTICALLY REGULATE NODULATION AND AFFECT ARBUSCULAR MYCORRHIZA IN *MEDICAGO TRUNCATULA*"

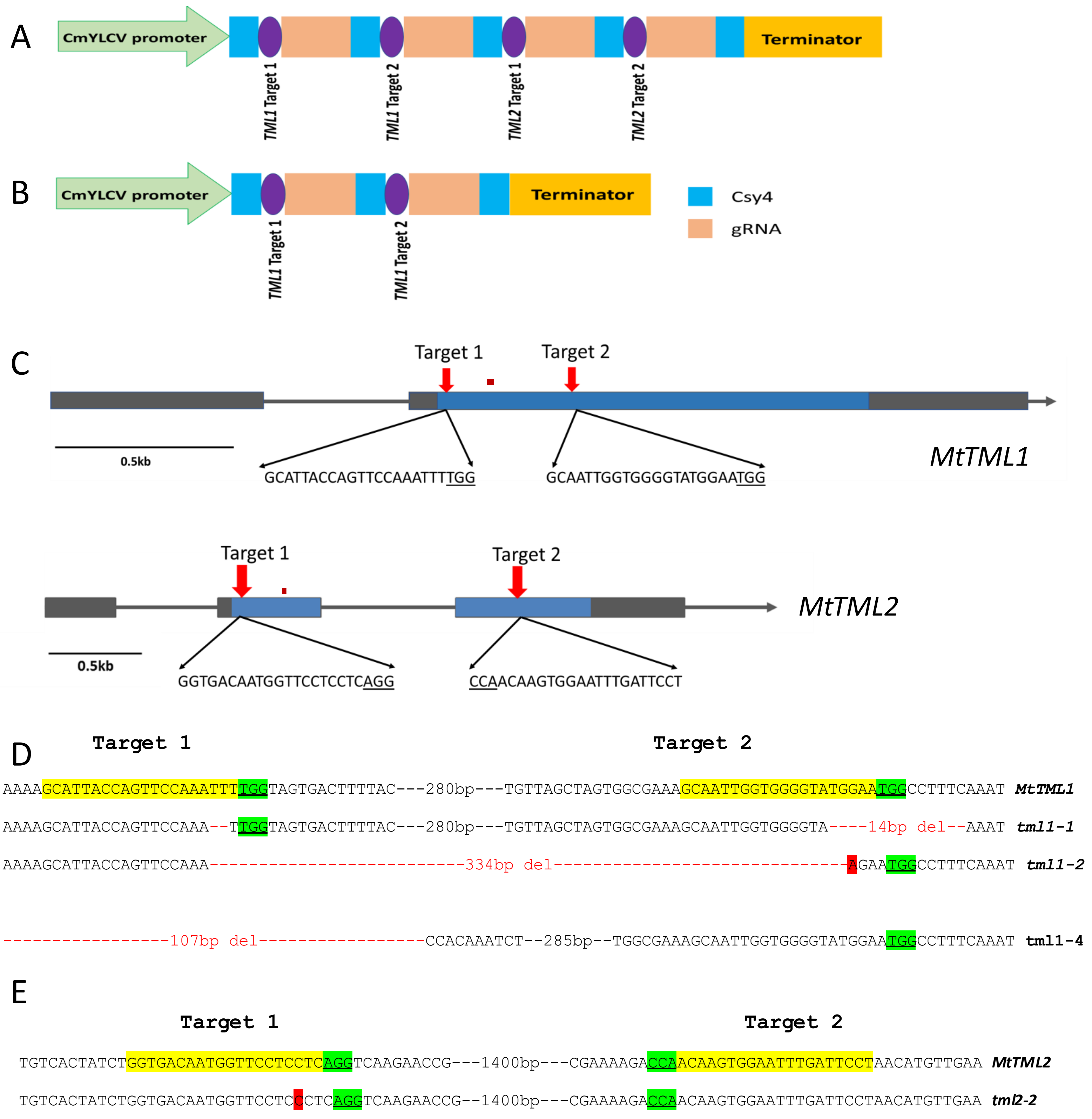


**Supplemental Figure 1. Multiplexed targeted CRISPR/Cas9 mutagenesis in *TML* genes using Csy4 plasmid**. (A) Schematic diagram of polycistronic gene construct used in pDIRECT_23C+TML1/2 for making *tml1/tml2* double mutants and (B) in pDIRECT_23C+TML1 for making *tml1* mutant (C) Maps of *MtTML1* and *MtTML2* genes showing the location of targets. PAM for each target is underlined. Blue boxes indicate CDS, grey boxes indicate UTR, red rectangles indicate predicted miRNA2111 binding sites, and lines indicate introns; arrow at the right end of grey box points to 3' end of gene. (D) CRISPR effects on DNA resulting in *tml1* alleles, displayed in relation to the target sequences (yellow) and PAM sequences (green). Base changes and deletions are in red. The *tml1-4* allele has a 107 bp deletion that removes 48bp upstream of the transcription start site and 59bp in the CDS, including target 1. (E) CRISPR effects on DNA in the *tml2* allele, displayed in relation to the target sequences (yellow) and PAM sequences (green). Base changes and deletions are in red.
