## Supplemental Figure 2 for "*TML1* AND *TML2* SYNERGISTICALLY REGULATE NODULATION AND AFFECT ARBUSCULAR MYCORRHIZA IN *MEDICAGO TRUNCATULA*"

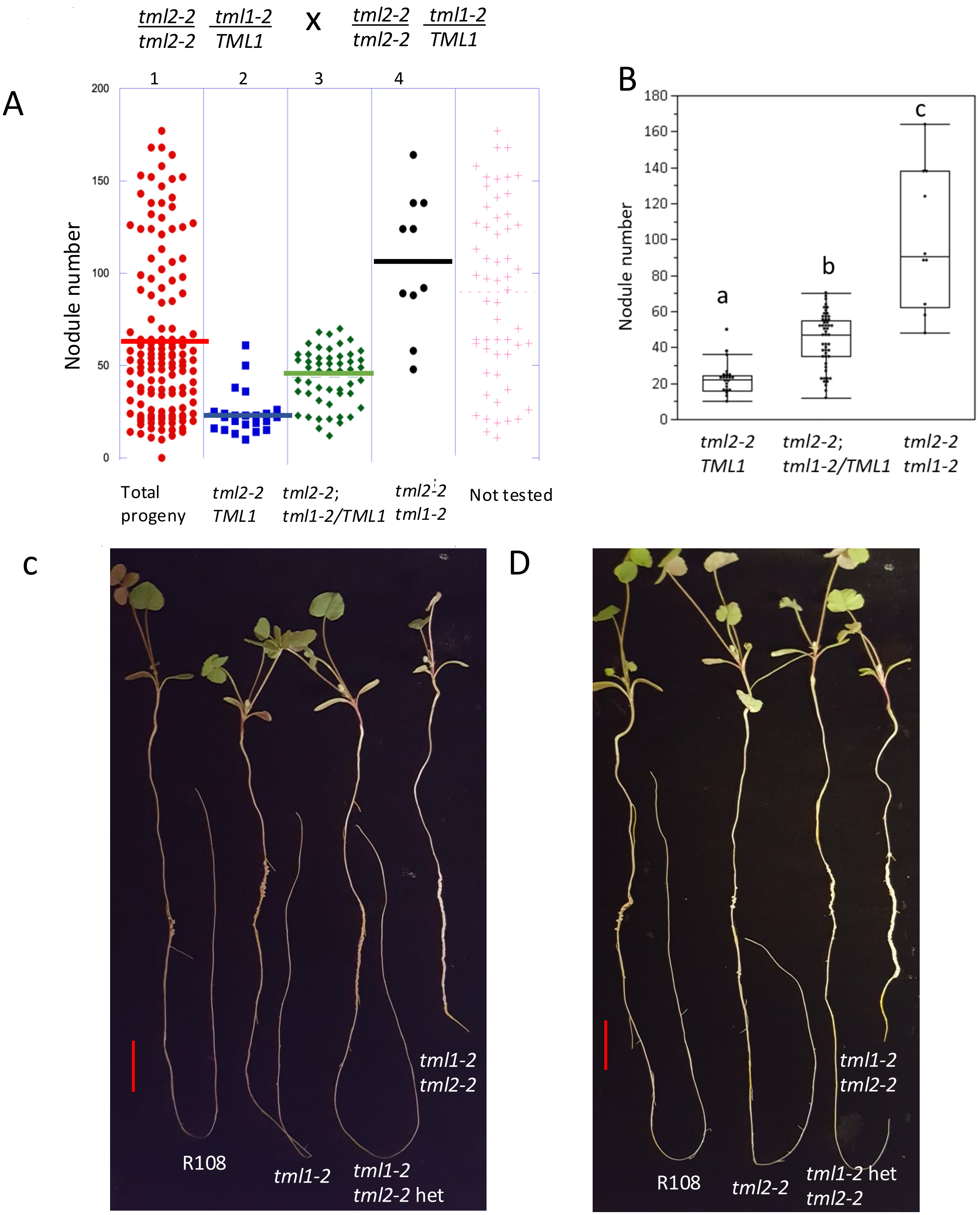


**Supplemental Figure 2. Molecular genotype analysis and photographs supporting heterozygous effect.** (A) Testing 60% (84) of the total population of 141 (indicated in red dots in panel 1) broken down in panels by molecular genotype determined by PCR. Blue dots (panel 2) are homozygous wild type *TML1,* green dots (panel 3) are heterozygous *tml1/TML1* and black dots (panel 4) are homozygous *tml1/tml1*. Lines indicate median nodule number for each group. (B) Box plot showing the distribution of nodule number in each genotype. Groups indicated by different letters are significantly different as tested by Tukey all pair test (p<0.001). (C) One representative plant per genotype L to R: R108 (wild type), *tml1-2*, *tml1-2 tml2-2* het, *tml1-2 tml2-2* and (D) R108 (wild type), *tml2-2*, *tml1-2* het *tml2-2*, and *tml1-2 tml2-2*. Scale bar = 2 cm.
