## Supplemental Table 2 for "*TML1* AND *TML2* SYNERGISTICALLY REGULATE NODULATION AND AFFECT ARBUSCULAR MYCORRHIZA IN *MEDICAGO TRUNCATULA*"

| **Primer number** | **Primer sequences (5’-3’)** | **Description** |
| --- | --- | --- |
|  | **Primers for *MtTML2 Tnt1* insert verification** |  |
| 2189 | ATTTGACTCTAATTTATTCTGTGA | Amplifies 1151 bp from WT *MtTML2*, Forward of 2189/2190 pair |
| 2190 | ATCTATGTTTGTCTTTTGCCAGTA | Reverse of 2189/2190 pair |
| 1925 | TCTGGATGAATGAGACTGGAGG | Tnt1 primer; Tnt1 insert in 5’ of *MtTML2* amplified by 2189/1925 |
| 2690 | CAGTGAACGAGCAGAACCTGTG | *Tnt1* primer; *Tnt1* insert in 3’ of *MtTML2* amplified and sequenced by 2690/3227; same primer pair was also used for insert verification in cDNA |
| 3227 | TTCCTTTCACCACCACTTGC | Positioned at exon 1 of *MtTML2*, Reverse |
| 2493 | TTGAAAATGGTGAAAAAGAGTCC | Positioned and exon 1 of *MtTML2* used for cDNA analysis, forward |
| 2494 | TCCACCAGCAACATAAGCAAAAT | Positioned at exon 2 of *MtTML2*, reverse for 2493 |
|  | **sgRNA primers for CRISPR construct** |  |
| 3079 | TCGTCTCCAACTGGTAATGCCTGCCTATACGGCAGTGAAC | CSY_MtTML1_crispr_Target 1 |
| 3080 | TCGTCTCAAGTTCCAAATTTGTTTTAGAGCTAGAAATAGC | REP_MtTML1_crispr_Target 2 |
| 3081 | TCGTCTCCCCCACCAATTGCCTGCCTATACGGCAGTGAAC | CSY_MtTML1_crispr_Target 2 |
| 3082 | TCGTCTCATGGGGTATGGAAGTTTTAGAGCTAGAAATAGC | REP_MtTML1_crispr_Target 2 |
| 3083 | TCGTCTCCACCATTGTCACCCTGCCTATACGGCAGTGAAC | CSY_MtTML2_crispr_Target 1 |
| 3084 | TCGTCTCATGGTTCCTCCTCGTTTTAGAGCTAGAAATAGC | REP_MtTML2_crispr_Target 1 |
| 3085 | TCGTCTCCAATTCCACTTGTCTGCCTATACGGCAGTGAAC | CSY_MtTML2_crispr_Target 2 |
| 3086 | TCGTCTCAAATTTGATTCCTGTTTTAGAGCTAGAAATAGC | REP_MtTML2_crispr_Target 2 |
| 3076 | TGCTCTTCGCGCTGGCAGACATACTGTCCCAC | CmYLCV (universal primer) (Cermak, et al., 2017) |
| 3077 | TGCTCTTCTGACCTGCCTATACGGCAGTGAAC | CSY_term (universal primer) (Cermak, et al., 2017) |
|  | **Primers for transgenic lines and CRISPR mutant analysis** |  |
| 3182 | CTGAGGCTACCAGACTCAAGA | *Cas9* specific primer forward |
| 3183 | AGGTTCTCAAGCCTTCTTGAC | *Cas9* specific primer reverse |
| 3065 | GATTATGAGCTGTTAAGAGGA | *MtTML1* mutation analysis forward (*tml 1-2* is 333bp shorter than WT; *tml1-3* BstXI digestion 105bp + 389 bp) |
| 3066 | TGTAATCTGATTGAATCCTTGG | *MtTML1* mutation analysis reverse |
| 3358 | AGTATCCGAATCATATCGCCA | *MtTML1* sequencing primer (paired with 3065 for analysis of *tml1-1* and *tml1-4.* In tml1-1, HaeIII digestion site lost – no digestion and *tml1-4* is 107bp shorter than WT) |
| 3148 | GAGTTATAGTGTGTCTAGTA | *MtTML2* mutation analysis forward; also used for *Tnt1* insert verification in *tml2-1* cDNA paired with 3149 and 1925 |
| 3149 | TTTCCCAAAGATTTGAATGCCA | *MtTML2* mutation analysis reverse |
|  | **Primers for Real Time PCR** |  |
| 2491 | GTGGCTGGGGTGTCGCATTTA | TML1 cDNA forward |
| 2492 | TCCCATCATCACAGCACAGTTCCT | TML1 cDNA reverse |
| 2493 | TTGAAAATGGTGAAAAAGAGTCC | TML2 cDNA forward |
| 2494 | TCCACCAGCAACATAAGCAAAAT | TML2 cDNA reverse |
| 2003 | GCAATGTGGGGATTTAGAGATT | NIN cDNA forward |
| 2004 | GGAAGATTGAGAGGGGAAGCTT | NIN cDNA reverse |
| 2450 | GCAGATAGACACGCTGGGA | PIK cDNA forward |
| 2451 | AACTCTTGGGCAGGCAATAA | PIK cDNA reverse |
